# The clarifying role of time series data in the population genetics of HIV

**DOI:** 10.1101/495275

**Authors:** Alison F. Feder, Pleuni S. Pennings, Dmitri A. Petrov

## Abstract

HIV can evolve remarkably quickly in response to anti-retroviral therapies and the immune system. This evolution stymies treatment effectiveness and prevents the development of an HIV vaccine. Consequently, there has been great interest in using population genetics to disentangle the forces that govern the HIV adaptive landscape (selection, drift, mutation, recombination). Traditional population genetics approaches look at the current state of genetic variation and infer the processes that can generate them [1, 2, 3, 4]. However, because HIV evolves rapidly, we can also sample populations repeatedly over time and watch evolution in action [5, 6, 7]. In this paper, we demonstrate how time series data can bound evolutionary parameters in a way that complements and informs traditional population genetic approaches.

Specifically, we focus on our recent paper [2], in which we show that, as improved HIV drugs have led to fewer patients failing therapy due to resistance evolution, less genetic diversity has been maintained following the fixation of drug resistance mutations. We interpret this as evidence that resistance to early HIV drugs that failed quickly and predictably was driven by soft sweeps while evolution of resistance to better drugs is both less frequent and when it takes place it is associated with harder sweeps due to an effectively lower HIV population mutation rate (*θ*). Recently, Harris et al. have proposed an alternative interpretation [8]: the signal could be due to an increase in the selective benefit of mutations conferring resistance to better drugs. Therefore, better drugs lead to faster sweeps with less opportunity for recombination to rescue diversity. In this paper, we use time series data to show that drug resistance evolution during ineffective treatment is very fast, providing new evidence that soft sweeps drove early HIV treatment failure.

## 1 Diversity signatures from a single time point provide information about evolution

In population genetics, we often sample data at a single time point and build models to explain how the observed patterns of genetic diversity might be generated. Many species evolve so slowly that researchers cannot directly observe it and working with snapshot data is the only possibility.

One particularly active field of study uses patterns of genetic diversity to understand the historical and ongoing role of selection. Selection can lead to many characteristic signatures, including the depression of diversity surrounding a recently fixed beneficial allele. As a single adaptive allele rises to high frequency, hitchhiking genetic neighbors also fix in the population in an event known as a **hard selective sweep.** The ratio of selection strength and recombination rate governs the distance on the chromosome from the adaptive site with depressed diversity following a sweep. The most widely used methods for detecting positive selection scan the genome for these sweep signatures [9, 10].

More recently, there has been increased attention to cases in which beneficial mutations enter the population rapidly and repeatedly. If a beneficial mutation lands on multiple genetic backgrounds before any single background can sweep, the backgrounds carrying the beneficial mutation will spread concurrently. In such a case, substantially more genetic diversity will be retained following the fixation of the beneficial mutation, because potentially diverse genetic backgrounds are brought to intermediate frequency. This is known as a **soft selective sweep.** Because hard and soft sweeps leave different diversity signatures, traditional selective scans do not detect them sensitively, and new approaches have been developed to identify soft sweeps [11, 12].

Whether sweeps will be hard or soft depends largely on the rate at which beneficial mutations enter the population each generation, which can be computed by the effective population size (*N_e_*) times the beneficial mutation rate (*μ_b_*). Note that the effective population size must be computed over the timescale relevant to adaptation [13, 14, 15]. The product of these terms (called *θ*) controls how long the population must wait to produce a beneficial mutation with fitness 1 + *s* relative to the non-mutant (≈ 1/*θ_s_* generations). If *θ* is small (*θ* ≪ 1), the adaptation is mutation-limited: in this regime, the population must wait between beneficial mutations, and when one does appear and establish, it will spread in the population to the exclusion of the ancestral type, in a hard sweep. However, if *θ* is large (*θ* ~ O(1)) and beneficial mutations enter the population at a rapid rate, adaptation is not mutation limited. Here the population does not need to wait long for a beneficial mutation to appear and others will appear after the first one. If one or more additional beneficial mutations enter the population before the first one fixes, the result is a soft sweep. It is therefore possible under certain circumstances to determine *θ* by looking for characteristic signatures of hard and soft sweeps [6, 15, 16].

Because of an interdependence through *θ*, whether a sweep will be hard or soft is also related to the probability of adaptation in a fixed period of time. Assuming that beneficial mutations sweep rapidly, when many mutations are expected to enter a population in quick succession (i.e., *θ* large), adaptation will be likely in a short period of time and sweeps will be soft. When the population must wait extensively for a beneficial mutation of large effect (θ small), adaptation is less likely to happen in a fixed period of time, and if it does happen, the sweep is more likely to be hard. Later in this paper, we will demonstrate this principle via simulations. However, the reasoning is much more general: if an independent event is likely to happen at least once, it is likely to happen multiple times.

## 2 Diversity signatures in single time point HIV data suggest two hypotheses characterizing early HIV evolution

As described above, we expect the probability of evolution in a fixed period of time to be dependent on *θ* and s. What might this imply about HIV evolution? Better HIV drugs have resulted in the probability of a patient evolving drug resistance in a year on therapy falling sharply from over 80% in the early 1990s to < 10% in the early 2010s. We might therefore expect that as the probability of resistance evolution has decreased, hard sweeps have become more prevalent. In [2] we examined differences in the diversity signatures of intra-patient HIV populations to understand if this decreasing prevalence of drug resistance was indeed accompanied by a transition from soft sweeps to hard sweeps, as might be expected if these better drugs had effectively increased the waiting time for a beneficial mutation. This is illustrated in the left-hand panels of Figure 1.

**Figure 1:**
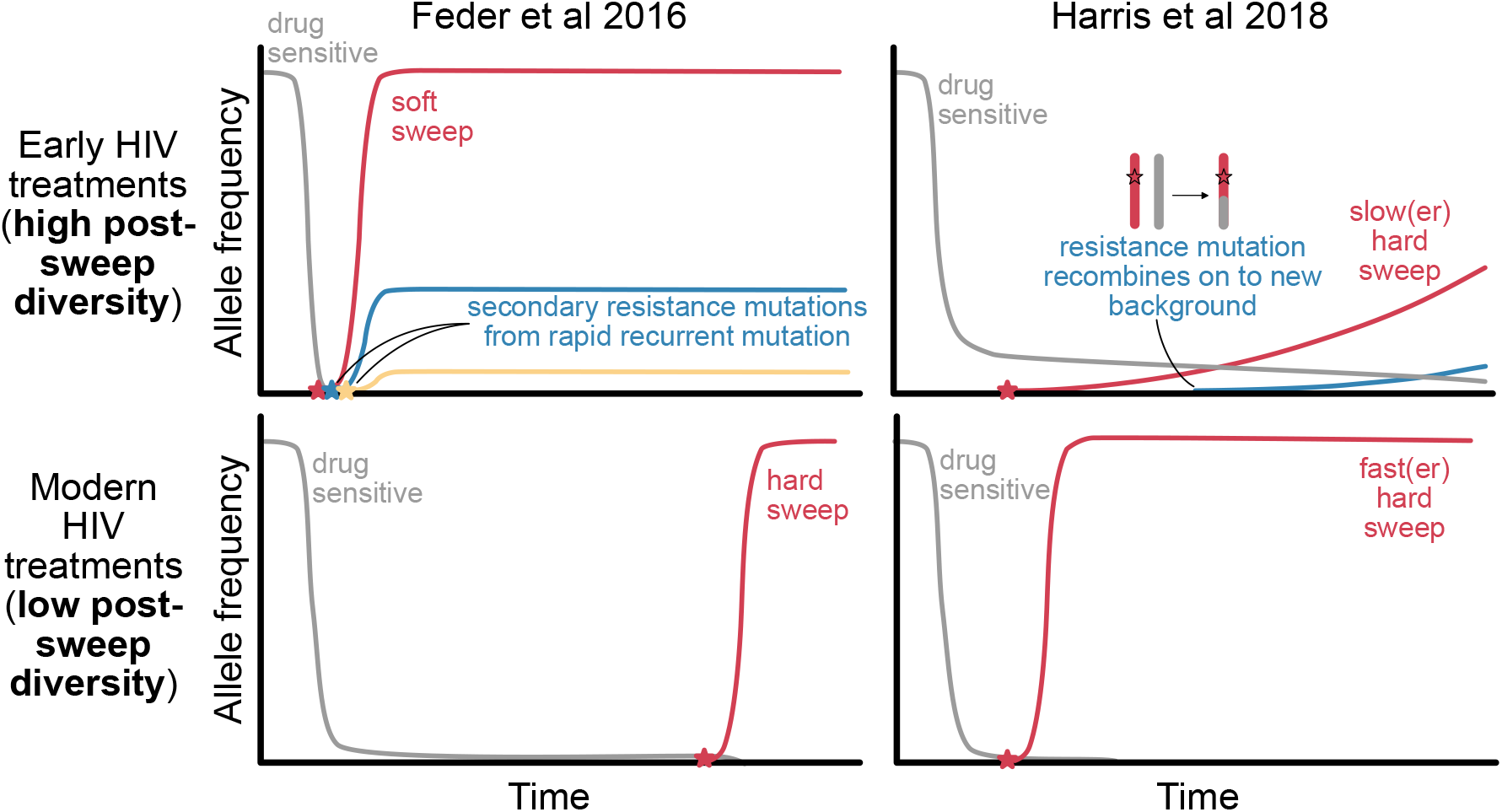
Cartoon of explanations for post-sweep diversity loss patterns observed in Feder et al. [2]. We seek to explain why more diversity is retained following the fixation of drug resistance to early, ineffective therapies than modern, effective ones. The **soft sweeps of recurrent mutations hypothesis** suggests that the primary difference is that *θ* is high for ineffective drugs, leading to short wait times and soft sweeps (top-left), whereas *θ* is lower for effective drugs, leading to long wait times and hard sweeps (bottom-left). The **recombination-rescued diversity hypothesis** suggests that the primary difference is the opportunity for recombination to rescue diversity following a beneficial mutation: sweeps in response to ineffective drugs are slow and occur after wider population bottlenecks, which gives recombination time to rescue diversity (top-right). Sweeps in response to effective drugs are fast and occur after narrow population bottlenecks, so recombination has less opportunity to rescue genetic diversity (bottom-right). Stars represent drug resistance mutations entering the population *de novo*, and drug pressure starts at the x-intercept.

Consistent with our hypothesis, we found that in response to early, ineffective therapies with a high probability of failure via drug resistance evolution, genetic diversity was not affected by the number of sweeps of resistance mutations within a patient. This is consistent with soft sweep signatures, whereby populations can fix mutations without losing genetic diversity. However, in response to good therapies with a low probability of resistance evolution, fixing a drug resistance mutation was associated with a significant loss in genetic diversity. This is consistent with the signature of a hard selective sweep.

Harris et al. [8] have recently proposed that different levels of recombination during a selective sweep can generate the patterns we observed in Feder et al. [2] without invoking soft sweeps. They argue that first, if ineffective drugs induce less severe population bottlenecks than those from effective drugs, more diversity will be present that can be rescued via recombination during a selective sweep. Second, if ineffective drugs have longer sweep times than effective drugs, recombination will have more time to recombine genetic diversity on to the sweeping haplotype. This hypothesis is illustrated in the right-hand panels of Figure 1.

Figure 1 shows that under both models, when resistance is acquired in response to poor drugs, little population diversity is lost. We discuss two explanatory hypotheses: soft sweeps in which recurrent mutation brings multiple genetic backgrounds to intermediate frequency, or slow sweeps that allow for the rescue of diversity via recombination (Upper row, Figure 1). These hypotheses cannot be readily distinguished using the snap shot data analyzed in Feder et al. [2]. However, because HIV is a well-studied system, we can draw upon external sources of information about drug resistance evolution in HIV to understand constraints on the timescales at play, specifically the speed and the probability of adaptation. We argue here that because time series data show that drug resistance evolution in HIV is fast and predictable, it is unlikely that this diversity is maintained through recombination-driven rescue of diversity.

## 3 HIV time series data suggest rapid adaptation and soft sweeps likely in early HIV evolution

Time series data show just how rapidly HIV can evolve. For example, Figure 2 shows the frequencies of drug resistance mutation M184V, which arises rapidly in 19 of 20 patients who were treated with 3TC monotherapy in the early 1990s [17]. Among the 20 patients, 16 had drug resistance over 50% frequency by week four (approximately 30 generations, at 1 HIV generation/day). Note, M184V is costly and unlikely to be present at high frequency as standing genetic variation before treatment [18]. Also note that four of the 20 patients had prior exposure to drugs for which M184V rendered partial resistance to therapy, but the percentage of these patients with M184V emerging within four weeks was very similar to the cohort as a whole (3/4 patients). This represents just one of many examples showing that HIV evolves drug resistance **rapidly** and **predictably** in response to early drug therapies [19, 20, 21]. These examples provide information about multiple population genetic parameters.

**Figure 2:**
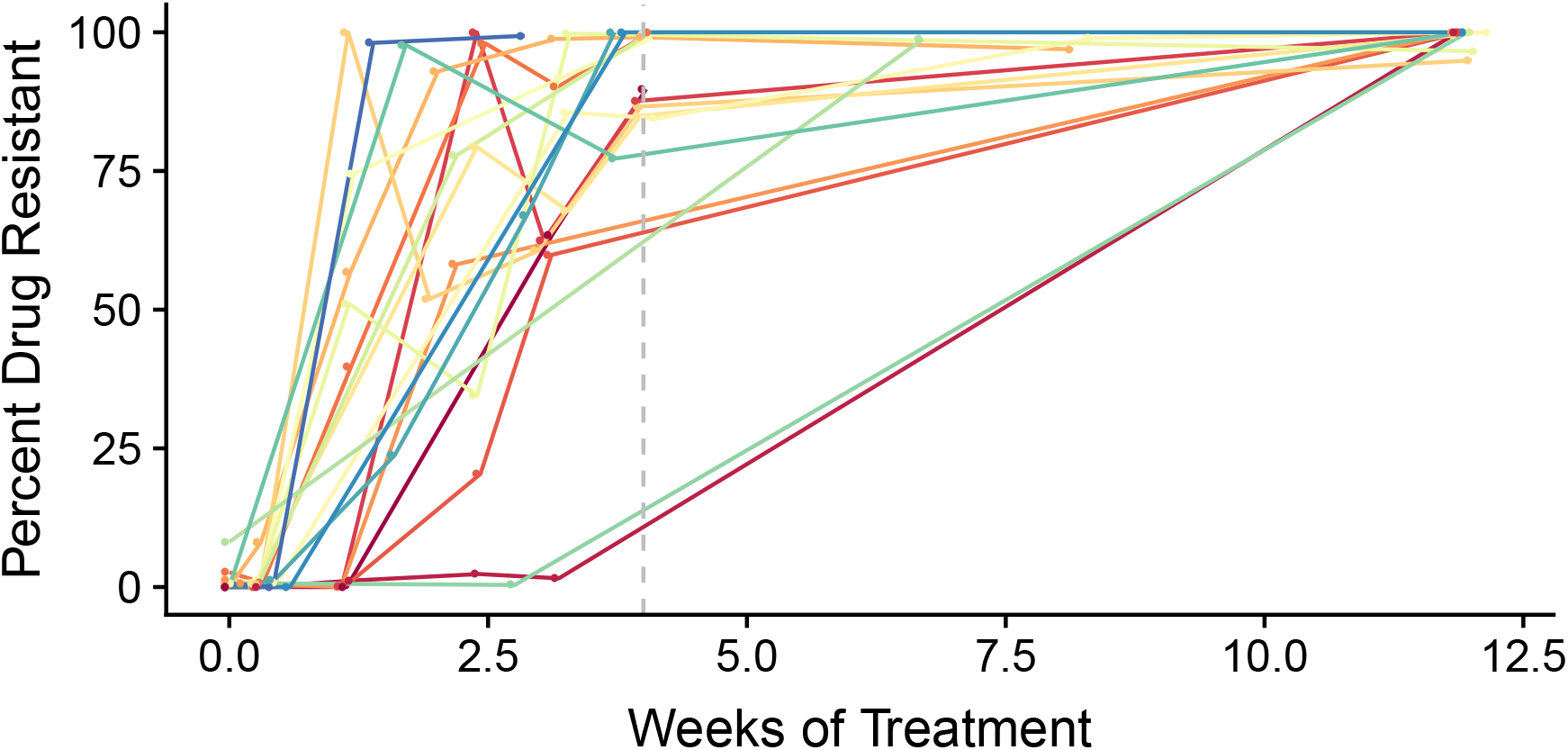
Frequencies of drug resistance mutation M184V in response to 3TC monotherapy in 20 patients. 16/20 have drug resistance observed above frequency 50% in 4 weeks (~ 28 generations, shown in dashed line). 19/20 have drug resistance above 50% in 12 weeks. The patient with no drug resistance is not plotted. Data replotted from Schuurman et al, 1995 [17] via WebPlotDigitizer [22].

HIV’s ability to adapt **rapidly** depends strongly on the strength of selection, s for the drug resistant variant under monotherapy. Understanding the sweep time of a beneficial variant (≈ log(*Ns*)/*s* generations when *s* ≪ 1) allows us to place bounds on *s*. The **predictability** of HIV’s evolution in a short period of time tells us about the number of beneficial mutations entering the population each generation, *θ*. That mutations appear promptly and consistently across populations suggests that *θ* must be high.

To quantify what the rapidity and predictably of drug resistance evolution can tell us about HIV evolution in response to monotherapy, we simulated a single locus model of drug resistance during evolution of resistance in HIV for a wide variety of values for *θ* and *s* (see Materials & Methods for full details). The value of *θ* is set by the mutation rate, which we set at 10^−5^ per site [23] and on the short-term effective population size relevant for such rapid adaptation. This short-term effective population size is bounded by the census population size of HIV, which can be on the order of 10^9^ and possibly much larger [24] (giving us *θ*10^4^). The effective population size is smaller than the census size, but how much smaller depends on many biological specifics of the HIV population in question, such as the proportion of infectious particles. Here, we varied *θ* from 10^−3^ to 10. For each parameter combination, we recorded the probability that beneficial mutations reached frequency ≥ 50% in 30 generations (Figure 3A), as an analog to the data presented in Figure 2. For sweeps to be very predictable on this timescale (80% of cases), both *s* and *θ* must be large: s needs to be on the order of 1 and *θ* must be greater than 0.1. For higher, potentially more realistic values of N (or, more correctly, the short-term *N_e_* relevant to rapid adaptation on the time scales of tens of generation) [6, 13], even higher s values are needed for sweeps as fast as observed.

**Figure 3:**
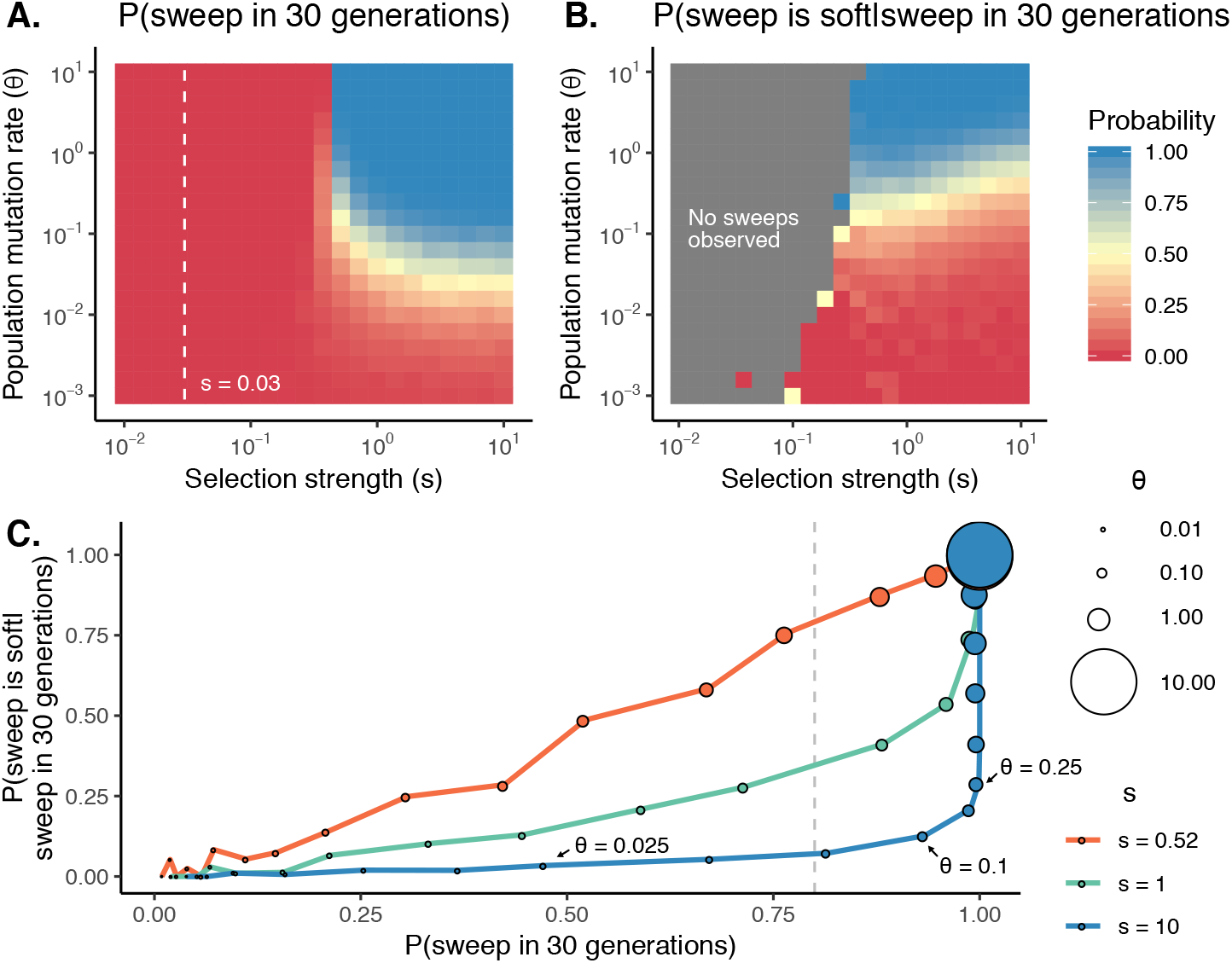
The probability of a sweep and a soft sweep are closely connected. For a variety of combinations of *θ* and *s*, we plot **A**) the probability of the one or more beneficial alleles reaching 50% frequency in 30 generations and **B**) we plot the proportion of sweeps that are soft conditional on occurring. Grey squares indicate the beneficial allele(s) did not reach 50% frequency in any simulations, and the white dashed line indicates the strongest selection strength simulated by Harris et al. (*s* = 0.03). In **C**), we plot the relationship between how likely a sweep is and its probability of being soft conditional on occurring and find that the two are strongly, positively correlated. *P*(*sweep*) = 80% (i.e., Schuurman 1995) is shown in a dashed line. Point size shows increasing values of *θ* for fixed values of *s* (*s* = (0.52,1,10), shown in red, green and blue). Other simulation parameters: 10^3^ replicates, *μ* = 10^−5^.

We then asked, conditional on a sweep occurring at each combination of parameters, what is the probability of that sweep being soft (Figure 3B). The parameter combinations in which sweeps are likely (i.e. > 80% probability of a sweep in 30 generations), are exactly those in which soft sweeps are most likely to occur. Specifically, as *θ* increases, the probability of both a sweep and a soft sweep increases (Figure 3C).

Note, the probability of a soft sweep depends on both *θ* and *s*, and if *s* ≫ *θ*, hard sweeps can occur because the sweep time is less than the waiting time for a new beneficial mutation. For example, when *s* = 10, *θ* = 0.1, sweeps are likely to occur in 30 generations (93%), but they are hard more often than not (87% of sweeps hard). However, even when s is so large, there is a relatively constrained part of the parameter space with respect to *θ* that makes both sweeps and hard sweeps very likely. Specifically, *θ* must be on order 0.1: indeed, if *θ* < 0.025, sweeps do not happen with necessity (< 50% frequency), but if *θ* > 0.25, sweeps are soft more than 25% of the time. Moreover, if *θ* ≈ 0.1 and s is large, these hard sweeps will retain little diversity anyway, and thus are unlikely to be the dominant mode of early adaptation given the observation of diversity retention in Feder et al. 2016 [2].

Note that standing genetic variation may also contribute to these sweeps. However, for these mutations to be reliably present before the start of treatment, *θ* must also be high. Although this does depend on the fitness cost of the mutation in the absence of the drug, much more important for predictably of presence is the population-wide mutation rate [25]. In the *θ* regimes in which the beneficial mutation is likely to arise via standing genetic variation, it is also likely to arise multiply via recurrent mutation after the onset of treatment [26].

The simulations above suggest that when sweeps happen on the timescales observed in early HIV drug resistance evolution data, we might expect them to be driven at least in part via soft sweeps of recurrent mutations. Indeed, we have clear, corroborating evidence that soft sweeps happen in HIV treated with ineffective therapies. Pennings, Kryazhimskiy and Wakeley studied how a strongly beneficial drug resistance mutation to protease inhibitors, K103N, emerges in viruses from treated patient sampled over time. K103N can be created by either an *A* → *T* or *A* → *C* mutation at the third nucleotide of the codon. Among a dataset of 17 patients fixing K103N in response to therapy, both *A* → *T* and *A* → *C* mutations were simultaneously observed in > 40% (7/17) of patients. One such patient is shown in Figure 4. At day 0, no drug resistance is present in 8 sequenced viral samples. At day 28, drug resistance is visible in two different encodings. Drug resistance must therefore have arisen from at least two independent mutations within these patients [6], and could not have been generated via recombination of a single mutation on to different backgrounds. This is a conservative estimate of the number of soft sweeps, because when two or more mutations of the same type happen (e.g., *A* → *T* occurs twice), the soft sweep appears to us as hard. There are other examples of soft sweeps in response to ineffective therapies from studies where multiple sequences are available per patient [19, 27].

**Figure 4:**
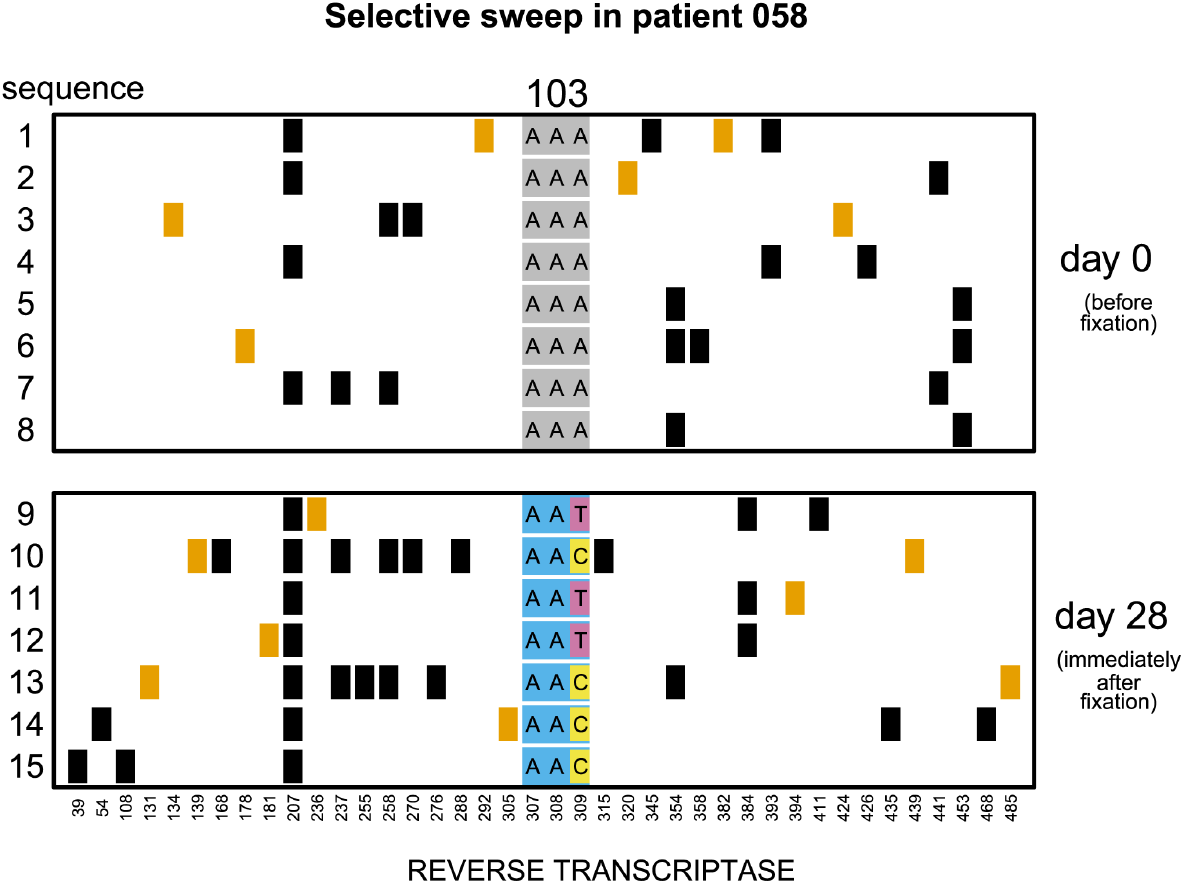
Soft selective sweep from Pennings, Kryazhimskiy and Wakeley [6]. Viruses were sampled from a patient before (day 0) and after (day 28) the fixation of drug resistance mutation K103N. At site 103, both *A* → *T* and *A* → *C* mutations spread at the third codon position. All genetically variable sites are plotted, with black squares representing synonymous substitutions and orange squares representing nonsynonymous substitutions.

## 4 Rapid adaptation allows recombination little opportunity to rescue diversity

Given the constraints on HIV’s evolutionary process in response to monotherapy demonstrated above, we return to the hypothesis that slow hard sweeps in response to ineffective drugs allow the rescue of genetic diversity. Recombination during a selective sweep rescues genetic diversity proportionally to the ratio of the selection strength and the recombination rate (*s/r*) [28]. The strength of selection therefore determines whether recombination will result in rescue of genetic diversity (assuming *r* fixed).

Harris et al. suggest that if sweeps in response to ineffective drugs are slow (*s* = 0.0075), they can fix mutations without a loss of diversity. They compare these sweeps to those in response to simulated modern drugs (*s* = 0.03) which do lose genetic diversity with the sequential fixations of drug resistance mutations [8]. However, time series data suggest that selection in response to monotherapy is stronger than either of these values (Figures 2 and 3). The selection strength simulating monotherapy (*s* = 0.0075) is at least two orders of magnitude too low to recapitulate the speed of drug resistance evolution expected from Figure 3. To contextualize, a sweep in response to ineffective drugs under this strength of selection would take *log*(*Ns*)/*s* ≈ 1.5 years, as opposed to the < 30 days suggested above (using *N* = 10^4^). Even the strongest selection strength considered (*s* = 0.03) is not strong enough to lead to any sweeps within four weeks. Since even stronger selection observed in data will lead to even more drastic reductions in diversity, the recombination-rescued diversity hypothesis cannot explain how diversity is maintained for HIV responding to ineffective drugs.

We also note that the recombination rate put forward in support of the recombination-rescued diversity hypothesis may be too high. It was estimated from untreated patients where coinfection (and therefore recombination) is more likely than in the simulated bottlenecked HIV populations [29]. Harris et al. therefore likely overestimate how much diversity will be maintained due to recombination during a sweep. Because HIV populations treated with monotherapies experience both much stronger selection and potentially weaker recombination than what was assumed by Harris et al. [8], abundant recombination during slow hard sweeps does not represent a viable alternative to create the patterns observed in Feder et al. [2].

## 5 Conclusion

Population genetics is a powerful tool to translate modern day genetic variation into an understanding of the forces acting over evolutionary history. Integrating time series data can provide important calibration on our understanding of the evolutionary process. For example, even small amounts of ancient DNA found in specific places at specific times have transformed the understanding our history as a species. In the case of HIV, detailed time series data help provide strong bounds on the population genetic forces at play. Through tracking changing populations over time, we can see that HIV has high *θ* and large *s* for drug resistance mutations in response to ineffective therapies.

Bounding the timescales of these population genetic forces can help us understand whether soft sweeps of multiple recurrent mutations or hard, slow sweeps with recombination can explain how HIV populations became resistant to early therapies without losing genetic diversity. When populations fix beneficial mutations rapidly and predictably, soft sweeps are not only possible, but probable. While slow, hard sweeps can retain diversity after a sweep in general, parameter constraints on *s* and *θ* suggest that this is not an appropriate model for HIV.

## 6 Materials & Methods

We simulated a one locus model of drug resistance with a fixed mutation rate of *μ* = 10^−5^ [23]. We simulated a range of parameter values with *N* = 10^2^ − 10^6^ (resulting in *Nμ* = *θ* = 0.001 −10) and *s* = 0.01 − 10. For each parameter combination, we ran 10^3^ forward simulation replicates, and recorded the percentage of runs in which the sum of all drug resistance mutations reached 50% frequency by generation 30, which we called a sweep. If a sweep occurred, we also recorded whether more than one mutation was at frequency above 5%, which we recorded as a soft selective sweep.

## 7 Acknowledgements

The authors gratefully acknowledge useful discussions with (alphabetically) Benjamin Good, Oskar Hallatschek, Arbel Harpak, Joachim Hermisson, Jona Kayser, Bernard Kim, Grant Kinsler, Noah Rosenberg, and the Petrov, Pennings and Hallatschek labs. The authors would also like to thank Andrew Sackman for pointing out an error in our implementation of the single locus model. This work was supported in part by the Miller Institute for Basic Research in Science at the University of California Berkeley (AFF).

## References

[1] Fabio Zanini and Richard A Neher. Quantifying selection against synonymous mutations in HIV-1 env evolution. Journal of virology, pages JVI-01529, 2013.

[2] Alison F Feder, Soo-Yon Rhee, Susan P Holmes, Robert W Shafer, Dmitri A Petrov, and Pleuni S Pennings. More effective drugs lead to harder selective sweeps in the evolution of drug resistance in HIV-1. Elife, 5:e10670, 2016.

[3] Fabio Zanini, Vadim Puller, Johanna Brodin, Jan Albert, and Richard A Neher. In vivo mutation rates and the landscape of fitness costs of HIV-1. Virus evolution, 3(1), 2017.

[4] Kristof Theys, Alison F Feder, Maoz Gelbart, Marion Hartl, Adi Stern, and Pleuni S Pennings. Within-patient mutation frequencies reveal fitness costs of CpG dinucleotides and drastic amino acid changes in HIV. PLoS genetics, 14(6):e1007420, 2018.

[5] Fabio Zanini, Johanna Brodin, Lina Thebo, Christa Lanz, Göran Bratt, Jan Albert, and Richard A Neher. Population genomics of intrapatient HIV-1 evolution. Elife, 4:e11282, 2015.

[6] Pleuni S Pennings, Sergey Kryazhimskiy, and John Wakeley. Loss and recovery of genetic diversity in adapting populations of HIV. PLoS genetics, 10(1):e1004000, 2014.

[7] RAJ Shankarappa, Joseph B Margolick, Stephen J Gange, Allen G Rodrigo, David Upchurch, Homayoon Farzadegan, Phalguni Gupta, Charles R Rinaldo, Gerald H Learn, XI He, et al. Consistent viral evolutionary changes associated with the progression of human immunodeficiency virus type 1 infection. Journal of virology, 73(12):10489–10502, 1999.

[8] Rebecca B. Harris, Andrew Sackman, and Jeffrey D. Jensen. On the unfounded enthusiasm for soft selective sweeps II: Examining recent evidence from humans, flies, and viruses. PLOS Genetics, 14(12):1–21, 12 2018.

[9] Benjamin F Voight, Sridhar Kudaravalli, Xiaoquan Wen, and Jonathan K Pritchard. A map of recent positive selection in the human genome. PLoS biology, 4(3):e72, 2006.

[10] Pardis C Sabeti, Patrick Varilly, Ben Fry, Jason Lohmueller, Elizabeth Hostetter, Chris Cotsapas, Xiaohui Xie, Elizabeth H Byrne, Steven A McCarroll, Rachelle Gaudet, et al. Genome-wide detection and characterization of positive selection in human populations. Nature, 449(7164):913, 2007.

[11] Nandita R Garud, Philipp W Messer, Erkan O Buzbas, and Dmitri A Petrov. Recent selective sweeps in North American Drosophila melanogaster show signatures of soft sweeps. PLoS genetics, 11(2):e1005004, 2015.

[12] Daniel R Schrider and Andrew D Kern. S/HIC: robust identification of soft and hard sweeps using machine learning. PLoS genetics, 12(3):e1005928, 2016.

[13] Talia Karasov, Philipp W Messer, and Dmitri A Petrov. Evidence that adaptation in Drosophila is not limited by mutation at single sites. PLoS genetics, 6(6):e1000924, 2010.

[14] Benjamin A Wilson, Dmitri A Petrov, and Philipp W Messer. Soft selective sweeps in complex demographic scenarios. Genetics, pages genetics–114, 2014.

[15] Bhavin S Khatri and Austin Burt. Robust estimation of recent effective population size from number of independent origins in soft sweeps. bioRxiv, page 472266, 2018.

[16] Timothy JC Anderson, Shalini Nair, Marina McDew-White, Ian H Cheeseman, Standwell Nkhoma, Fatma Bilgic, Rose McGready, Elizabeth Ashley, Aung Pyae Phyo, Nicholas J White, et al. Population parameters underlying an ongoing soft sweep in southeast Asian malaria parasites. Molecular biology and evolution, 34(1):131–144, 2016.

[17] Rob Schuurman, Monique Nijhuis, Remko van Leeuwen, Pauline Schipper, Dorien de Jong, Phil Collis, Sven A Danner, John Mulder, Clive Loveday, Cindy Christopherson, et al. Rapid changes in human immunodeficiency virus type 1 RNA load and appearance of drug-resistant virus populations in persons treated with lamivudine (3TC). Journal of Infectious Diseases, 171(6):1411–1419, 1995.

[18] Vivek Jain, Maria C Sucupira, Peter Bacchetti, Wendy Hartogensis, Ricardo S Diaz, Esper G Kallas, Luiz M Janini, Teri Liegler, Christopher D Pilcher, Robert M Grant, et al. Differential persistence of transmitted HIV-1 drug resistance mutation classes. Journal of Infectious Diseases, 203(8):1174–1181, 2011.

[19] Paul Kellam, Charles AB Boucher, Jolanda MGH Tijnagel, and Brendan A Larder. Zidovudine treatment results in the selection of human immunodeficiency virus type 1 variants whose genotypes confer increasing levels of drug resistance. Journal of General Virology, 75(2):341–351, 1994.

[20] Brendan A Larder, Graham Darby, and Douglas D Richman. HIV with reduced sensitivity to zidovudine (AZT) isolated during prolonged therapy. Science, 243(4899):1731–1734, 1989.

[21] Charles AB Boucher, Eithne O’Sullivan, Jan W Mulder, Chitra Ramautarsing, Paul Kellam, Graham Darby, Joep MA Lange, Jaap Goudsmit, and Brendan A Larder. Ordered appearance of zidovudine resistance mutations during treatment of 18 human immunodeficiency virus-positive subjects. Journal of Infectious Diseases, 165(1):105–110, 1992.

[22] Ankit Rohatgi. WebPlotDigitizer, 2011.

[23] Michael E Abram, Andrea L Ferris, Wei Shao, W Gregory Alvord, and Stephen H Hughes. Nature, position, and frequency of mutations made in a single cycle of HIV-1 replication. Journal of virology, 84(19):9864–9878, 2010.

[24] Subgroup ‘Assessment of Pathogens Transmissible by Blood’ German Advisory Committee Blood (Arbeitskreis Blut). Human Immunodeficiency Virus (HIV). Transfusion medicine and hemotherapy: offizielles Organ der Deutschen Gesellschaft fur Transfusionsmedizin und Immunhamatologie, 43(3):203–222, 05 2016.

[25] Joachim Hermisson and Pleuni S Pennings. Soft sweeps and beyond: understanding the patterns and probabilities of selection footprints under rapid adaptation. Methods in Ecology and Evolution, 8(6):700–716, 2017.

[26] Nick Barton. Understanding adaptation in large populations. PLoS genetics, 6(6):e1000987, 2010.

[27] Alison F Feder, Christopher Kline, Patricia Polacino, Mackenzie Cottrell, Angela DM Kashuba, Brandon F Keele, Shiu-Lok Hu, Dmitri A Petrov, Pleuni S Pennings, and Zandrea Ambrose. A spatio-temporal assessment of simian/human immunodeficiency virus (SHIV) evolution reveals a highly dynamic process within the host. PLoS pathogens, 13(5):e1006358, 2017.

[28] John H Gillespie. Population genetics: a concise guide. JHU Press, 2010.

[29] Richard A. Neher and Thomas Leitner. Recombination Rate and Selection Strength in HIV Intra-patient Evolution. PLOS Computational Biology, 6(1):1–7, 01 2010.

